# OASIS: in vivo AAV-mediated transduction and genome editing of adult oligodendrocytes

**DOI:** 10.1101/2025.10.20.683551

**Authors:** Xiaoyun Ding, Jackson R. Curtis, Yueheng Xing, Yu Wu, Elior Peles, Matthew N. Rasband

**Author notes:** Correspondence should be addressed to M.N.R.

## Abstract

New viral approaches have revolutionized neuroscience by precisely delivering genes in neurons; for example, to control or monitor activity in specific neuronal cell types. In contrast, the manipulation of oligodendrocytes requires the Cre-LoxP system and gene-by-gene engineering, breeding, and genotyping. Here we introduce OASIS (<u>O</u>ligodendrocyte <u>A</u>AV-CRISPR mediated <u>S</u>pecific In vivo editing <u>S</u>ystem), a versatile platform that combines SELECTIV, an AAV-receptor-based transduction strategy, with HiUGE, an NHEJ-mediated CRISPR/Cas9 knock-in approach. We show efficient and specific oligodendrocyte transduction across the brain and tagging of endogenous cytoskeletal, myelin, cell adhesion, scaffolding, and junctional proteins. OASIS enables sparse yet reliable labeling, allowing direct visualization of a protein’s subcellular localization with single-cell resolution. We successfully fused the biotin-ligase TurboID with endogenous oligodendroglial Neurofascin-155, thereby achieving targeted biotinylation of the axoglial junction. OASIS is rapidly customizable for any gene-of-interest. Together, OASIS overcomes longstanding barriers in oligodendrocyte biology, providing a powerful system for precise, customizable genome editing and subcellular visualization in the adult brain.

## Introduction

A comprehensive understanding of a novel gene’s function in the nervous system requires cell type-specific gain- and loss-of-function approaches as well as precise expression profiling *in vivo*. Traditionally, this relies on the Cre-LoxP system and antibody-based immunofluorescence. However, generating new mouse lines is labor-intensive and time-consuming, and antibody-based approaches are often constrained by poor specificity, batch variability, or lack of availability. Unvalidated antibodies have become a leading contributor to the reproducibility crisis in biomedical research ^1^, underscoring the need for alternatives that allow direct labeling and visualization of endogenous proteins in vivo. For protein visualization, CRISPR/Cas9-based genome editing offers one solution by enabling the introduction of epitope tags into endogenous loci. Strategies such as Slender, ORANGE, HITI, and HiUGE use Adeno-Associated Virus (AAV) and homology directed repair (HDR) or non-homologous end joining (NHEJ) to achieve knock-in with varying efficiency ^2-4^. NHEJ-based HiUGE has been successfully applied in neurons to validate novel proteins identified through proteomics ^5-7^. In contrast to neurons and astrocytes, oligodendrocytes are highly resistant to AAV transduction. Thus, the application of these methods to oligodendrocyte-lineage cells remains limited by poor viral tropism and represents a significant barrier in myelin biology.

Glial biologists have often turned to in vitro culture systems, where oligodendrocyte precursor cells can be infected with retroviruses or lentiviruses to overcome these barriers. However, these models lack the extracellular matrix and axon-glia interactions that shape oligodendrocyte function *in vivo*, limiting their physiological relevance. Thus, a method to rapidly and efficiently manipulate oligodendrocytes in the intact brain is urgently needed.

Recently, the *SELECTIV* (<u>SEL</u>ective <u>E</u>xpression and <u>C</u>ontrolled <u>T</u>ransduction *In <u>V</u>ivo*) system was developed to overcome barriers in AAV delivery by overexpressing viral receptors in a cell type-specific manner, enabling efficient transduction of otherwise resistant cell types such as muscle stem cells ^8^; in addition, the *SELECTIV* mice also express spCas9 in the same cells expressing the AAV receptor. We reasoned that by coupling *SELECTIV* with oligodendrocyte-lineage-specific Cre drivers, robust tropism could be created in oligodendrocytes that captures AAV particles normally restricted to neurons or astrocytes; the presence of spCas9 could also be exploited for efficient genome editing. Here, we combined the strengths of *SELECTIV* and HiUGE to develop OASIS (<u>O</u>ligodendrocyte <u>A</u>AV-CRISPR mediated <u>S</u>pecific In vivo editing <u>S</u>ystem), a versatile platform that enables efficient transduction of oligodendrocytes for transgene expression or for precise *in vivo* gene editing. We demonstrate use of OASIS in adult mice for knock-in of tags for subcellular, single-cell resolution, and even enzymes for biochemical manipulation of local protein environments.

## Results

### Cell-type specific AAV receptor overexpression redirects AAV tropism toward oligodendrocyte lineage cells *in vivo*

The natural tropism of AAV towards neurons and astrocytes has been widely engineered and exploited in gene therapy. However, efficient transduction of oligodendrocyte (OL) lineage cells has remained challenging. Inspired by the *SELECTIV* system developed by Zengel et al. (2023) ^8^ which enables robust transduction of refractory cell types by AAV receptor overexpression (e.g., muscle stem cells), we reasoned that a similar artificial and increased tropism could be leveraged in OL-lineage cells in the central nervous system by overexpressing the AAV receptor (**Fig. 1A**, referred to as *OL-SELECTIV*), thereby reducing neuronal transduction while enhancing OL targeting.

**Figure 1.**
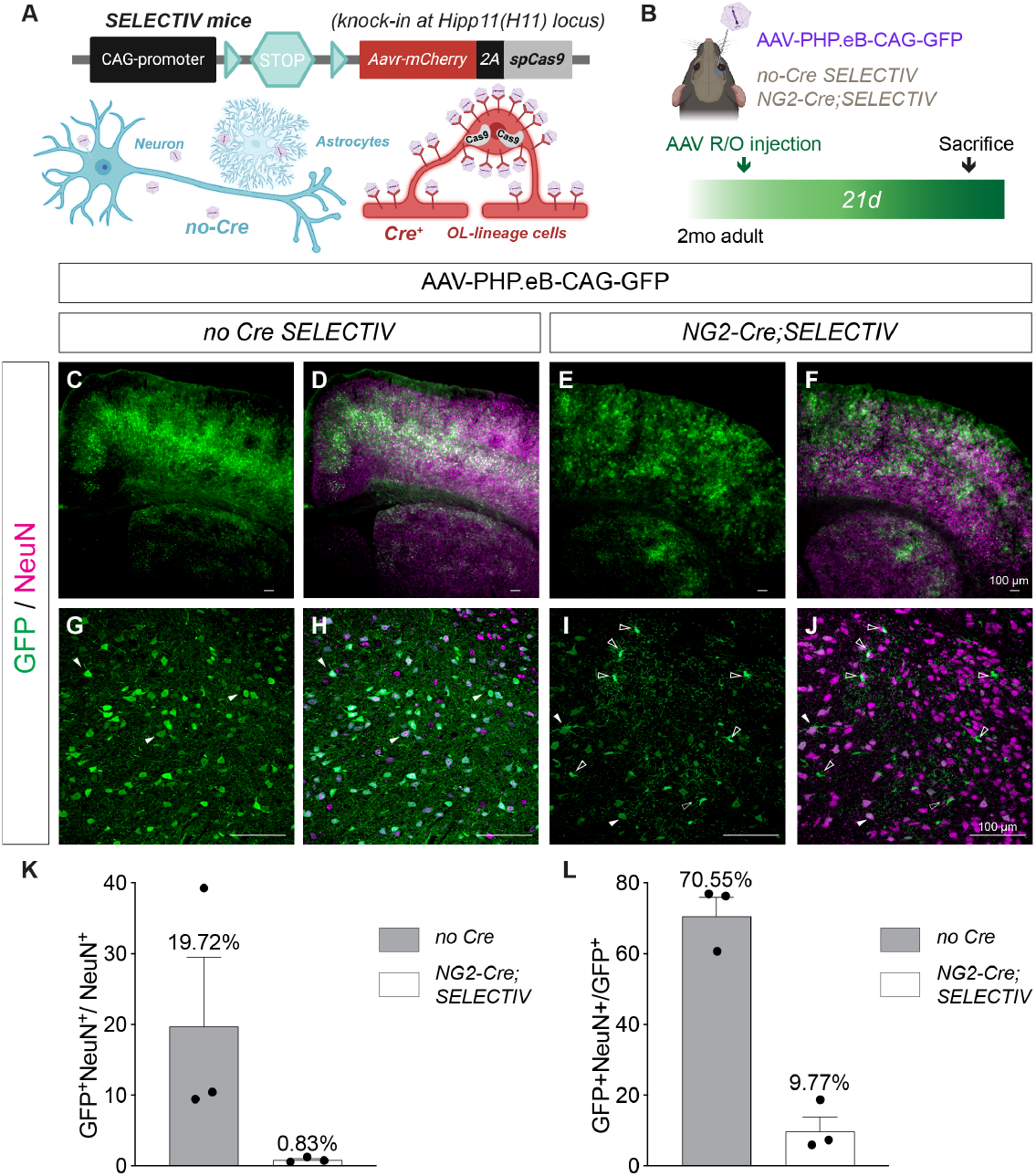
Cell-type specific AAV receptor overexpression redirects AAV tropism toward oligodendrocyte lineage cells in vivo. (A) Schematic of the SELECTIV system adapted for oligodendrocyte-lineage targeting. Cre-dependent overexpression of AAV receptors on the surface of oligodendrocyte-lineage cells increases AAV binding to oligodendrocytes and markedly reduces AAV tropism for neurons and astrocytes. (B) Experimental paradigm: AAVs were systemically delivered via retro-orbital (R/O) injection into 2-month-old adult control or *NG2-Cre;SELECTIV* mice and analyzed after 3 weeks. (C-F) Representative immunofluorescence co-labeling of GFP and NeuN in control and NG2-Cre SELECTIV mice. Scale bar, 100 μm. (G-J) Higher-magnification images from control and *NG2-Cre;SELECTIV* mice. Solid arrowheads indicate colocalization, empty arrowheads indicate absence of colocalization. Scale bar, 100 μm. (K) Quantification of GFP^+^NeuN^+^ cells among total NeuN^+^ cells. N = 3 mice per genotype. Control: 19.72 ±9.80% (mean ±SEM); *NG2-Cre;SELECTIV*: 0.84 ±0.20% (mean ±SEM). (L) Quantification of GFP^+^NeuN^+^ cells among total GFP^+^ cells. N = 3 mice per genotype. Control: 70.55 ±5.33% (mean ±SEM); *NG2-Cre;SELECTIV*: 9.77 ±4.07% (mean ±SEM).

To test this, we systemically delivered AAVPHP.eB-CAG-GFP by retro-orbital (R/O) injection and compared transduction in control *SELECTIV* versus *NG2-Cre;SELECTIV* mice after three weeks of incubation (**Fig. 1B**). In controls, GFP+ cells were predominantly neurons with axonal and dendritic morphology (**Fig. 1C-D, G-H**). In *NG2-Cre;SELECTIV* mice, neuronal labeling was markedly reduced and GFP+ cells exhibited typical OPC-like morphology instead (**Fig. 1E-F, I-J**). Quantification confirmed an over 20-fold decrease in the percentage of total neurons transduced by AAV (19.72 ±9.80% vs. 0.84 ±0.20%, N=3 per genotype; **Fig. 1K**). Nevertheless, ∼0.5-1% of neurons remained transduced, indicating that further optimization is required for exclusive OL targeting. And among all the transduced GFP+ cells, the percentage of neurons was drastically reduced by about 8-fold in *NG2-Cre;SELECTIV* mice compared to control (70.55 ±5.33% vs. 9.77 ±4.07%, N=3 per genotype; **Fig. 1L**).

### Different Cre drivers and viral capsids yield robust and stage-specific OL transduction

We next determined if *OL-SELECTIV* is broadly compatible with multiple Cre drivers and viral serotypes, and whether specific pairings maximize transduction. To further minimize neuronal expression, we used a Cre-dependent reporter (AAV-CAG-flex-GFP). We tested three different Cre drivers spanning OL development: *NG2-Cre* (for OPCs), *MOBP-iCreER*^*T2*^ (inducible, for pre-myelinating OLs), and *MOBP-iCre* (for mature OLs) (**Fig. 2A**). For AAV serotypes, we chose AAV-PHP.eB and AAVCAP-B10, due to their high efficiency in crossing the blood-brainbarrier and superior CNS targeting ^9-11^. We observed robust OL transduction across all six conditions (**Fig. 2B-I**).

**Figure 2.**
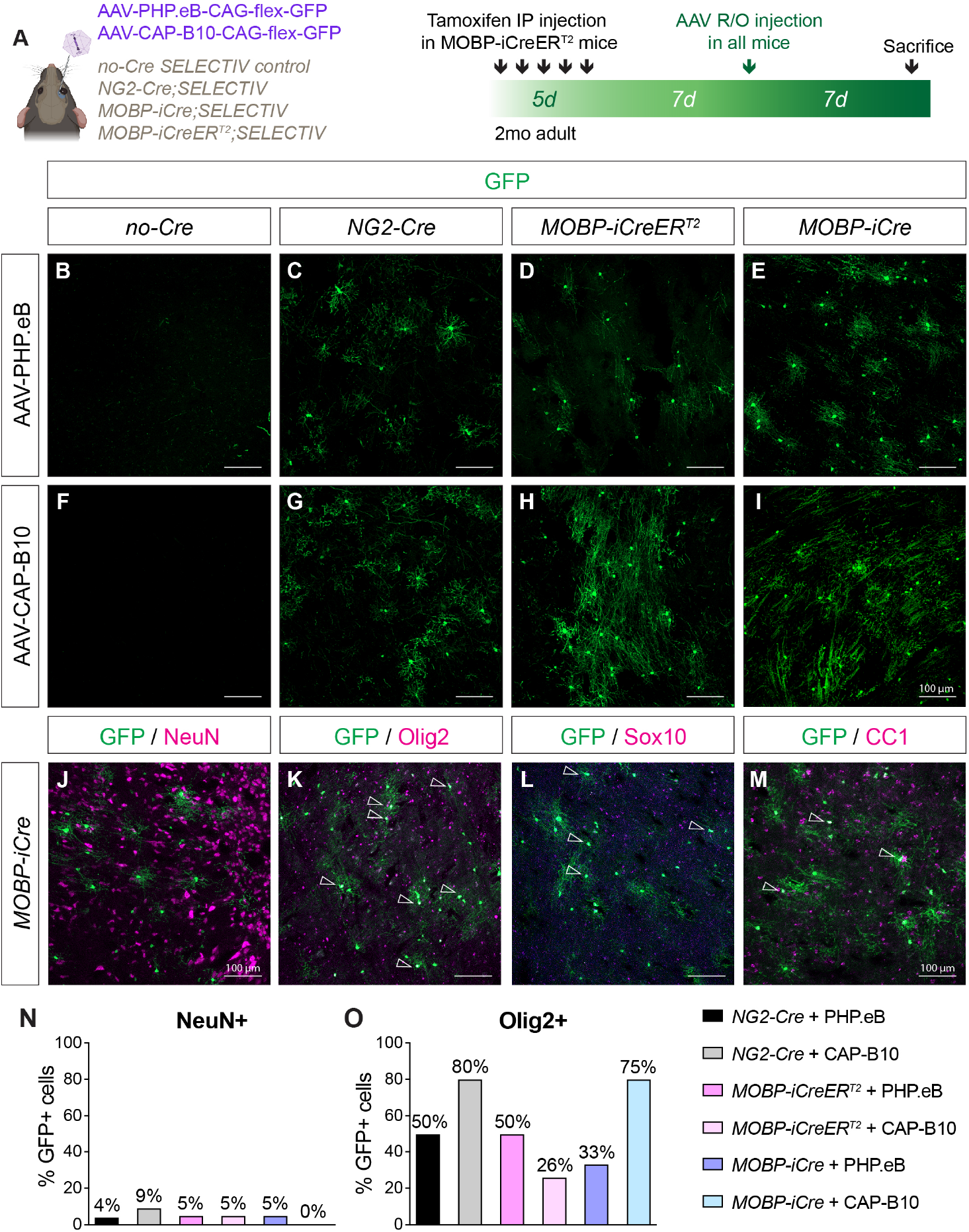
Different Cre drivers and viral capsids yield robust and stage-specific OL transduction. (A) Experimental paradigm: In 2-monthold *MOBP-iCreER*^*T2*^*;SELECTIV* mice, tamoxifen was delivered intraperitoneally (IP) for 5 consecutive days, followed by a 7-day incubation period prior to AAV administration. Control *SELECTIV, NG2-Cre;SELECTIV* and *MOBP-iCre;SELECTIV* mice were directly injected with AAV without tamoxifen induction. Two AAV serotypes (PHP.eB and CAP-B10) were systemically delivered via retro-orbital (R/O) injection to mice of four different genotypes and analyzed after 7 days. (B-E) Representative images showing GFP expression in the brain following AAV-PHP.eB administration in control (B), *NG2-Cre;SELECTIV* (C), *MOBP-iCreER*^*T2*^*;SELECTIV* (D), and *MOBP-iCre;SELECTIV* (E) mice. Scale bar, 100 μm. (F-I) Representative images showing GFP expression in the brain following AAV-CAP-B10 administration in control (F), *NG2-Cre;SELECTIV* (G), *MOBP-iCreER*^*T2*^*;SELECTIV* (H), and *MOBP-iCre;SELECTIV* (I) mice. Scale bar, 100 μm. (J-M) Lineage specific analysis of the GFP+ cells transduced by AAV-CAP-B10 in *MOBP-iCre;SELECTIV* mice. GFP expression was co-labeled with NeuN (neurons; J), Olig2 (oligodendrocyte lineage; K), Sox10 (oligodendrocyte lineage; L), and CC1 (mature oligodendrocytes; M). Empty arrowheads indicate colocalization. Scale bar, 100 μm. (N-O) Quantification of GFP+ cell identity, showing the percentage of NeuN^+^ cells (N) and Olig2^+^ cells (O) among total GFP^+^ cells.

The predominant cell state reflected the Cre driver used: *NG2-Cre* mice yielded GFP+ OPC-like cells (Fig. 2C, G), *MOBP-iCreER*^*T2*^ mice produced intermediate pre-myelinating OLs (Fig. 2D, H), and *MOBP-iCre* mice labeled mature myelinating OLs with more elaborate sheaths (Fig. 2E, I). Marker analysis confirmed minimal colocalization with NeuN (0-9%; Fig. 2J, N), while the majority of GFP+ cells expressed OL markers Olig2 (26-80%; Fig. 2K, O), Sox10 (Fig. 2L), and CC1 (Fig. 2M), consistent with OL lineage specificity.

**Figure 3.**
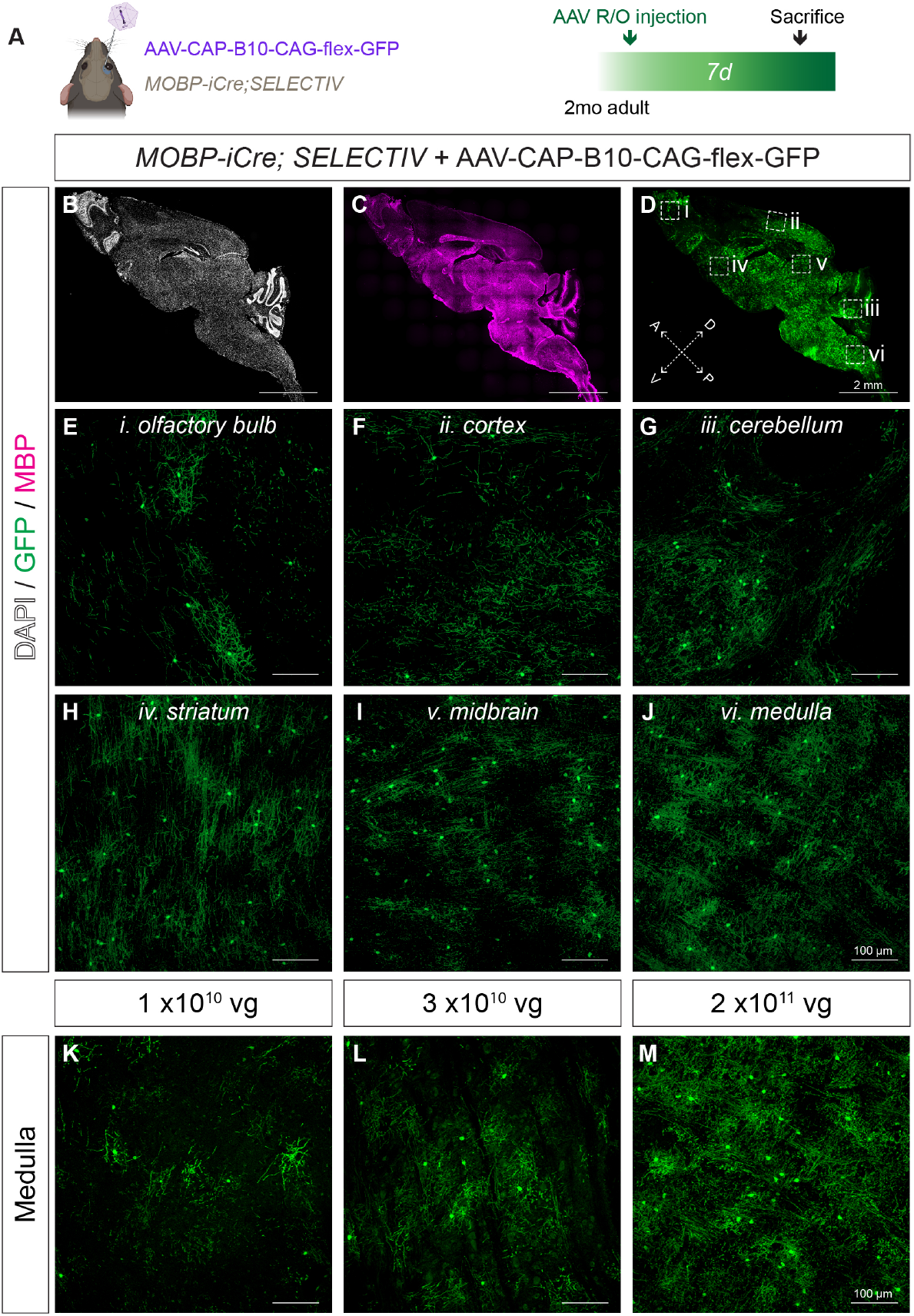
Oligodendrocyte transduction is efficient across brain regions and tunable by viral dosage. (A) Experimental paradigm: AAV-CAP-B10-CAG-flex-GFP was systemically delivered via retro-orbital (R/O) injection into 2-month-old adult *MOBP-iCre;SELECTIV* mice and analyzed after 7 days. (B-D) Representative sagittal brain sections showing whole-brain transduction. Nuclei were labeled with DAPI (B), white matter was marked by MBP immunostaining (C), and GFP expression was broadly detected across the brain (D). Six regions (i-vi) were further examined at higher magnification. A, anterior; P, posterior; D, dorsal; V, ventral. Scale bar, 2 mm. (E-J) Higher-magnification images from selected brain regions, including olfactory bulb (E), cortex (F), cerebellum (G), striatum (H), midbrain (I), and medulla (J). Scale bar, 100 μm.

### OL transduction is efficient across diverse brain regions and tunable by viral dosage

Because *MOBP-iCre;SELECTIV* mice injected with AAVCAP-B10-flex-GFP exhibited the highest OL-lineage specificity with minimal neuronal labeling, we examined regional transduction efficiency across the adult brain. Within seven days of injection, widespread GFP expression was observed in the olfactory bulb, cortex, cerebellum, striatum, midbrain, and medulla (**Fig. 3B-J**) of 2 month-old adult mice. Notably, the transduction efficiency was particularly high in the midbrain and medulla, where a high density of white matter is located. Moreover, increasing viral titer (from 1 × 10^10^ to 2 × 10^11^ vg/animal) significantly increased the number of transduced cells (**Fig. 3K-M**), demonstrating that OL transduction can be further fine-tuned by AAV dosage. These findings demonstrate that AAV receptor overexpression in OLs, combined with Cre-dependent expression, enables both specific and efficient OL transduction in vivo.

### OASIS enables endogenous protein tagging and subcellular visualization in OLs

Having established that the *OL-SELECTIV* system enables specific and efficient OL transduction, we next leveraged the spCas9 enzyme intrinsically expressed in these mice to perform in vivo genome editing of adult oligodendrocytes. Because knock-in (KI) efficiency is inherently lower than knockout, we first tested the most stringent application—tagging with spaghetti monster-V5 (smV5). We previously validated this design for tagging axon initial segment proteins in neurons ^6,7^ (**Fig. 4A-B**).

**Figure 4.**
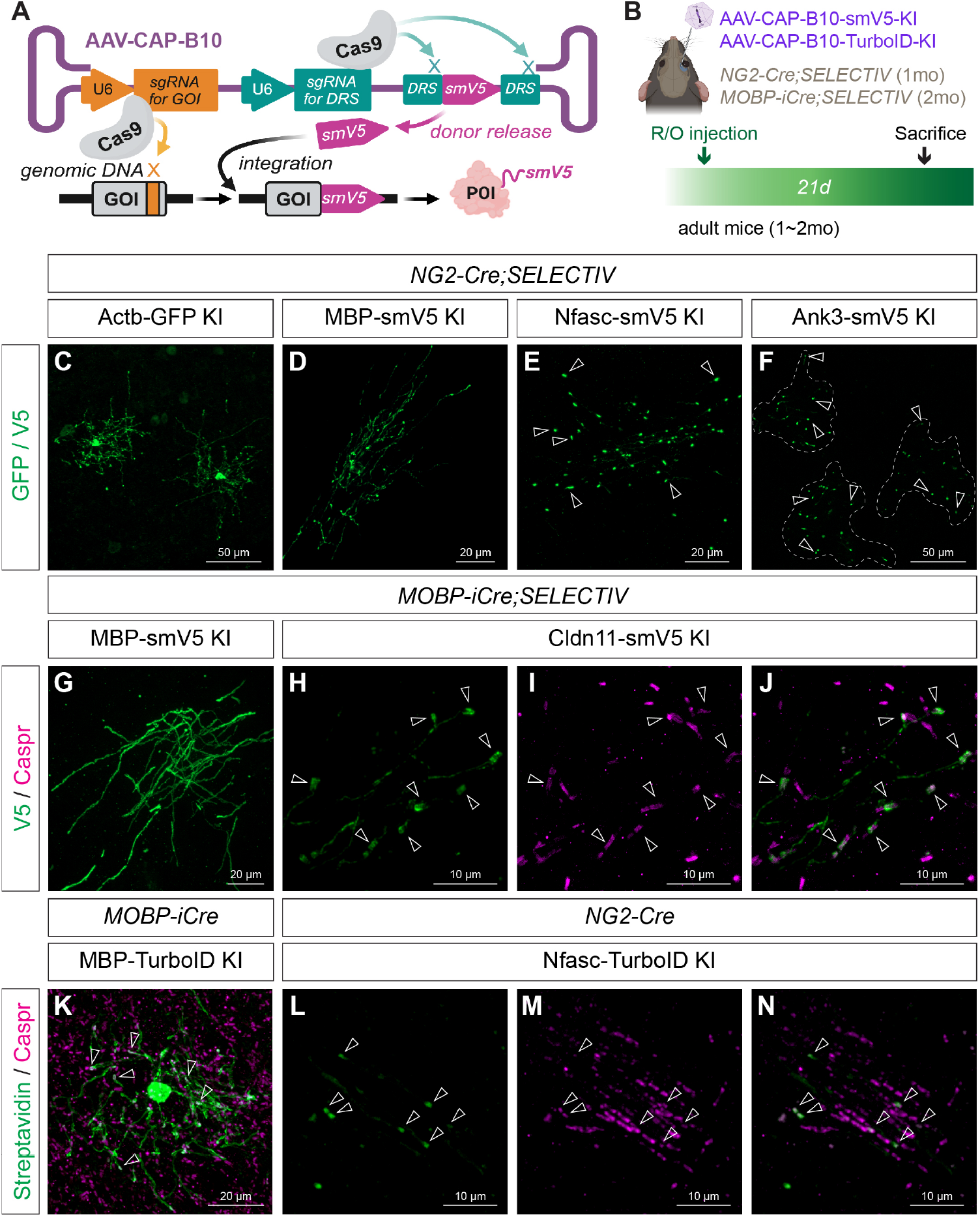
OASIS enables endogenogenous protein tagging and subcellular visualization in OLs. (A) Schematic of the NHEJ-based KI strategy. The construct carries both a genomic DNA-targeting sgRNA sequence for the gene-of-interest (GOI) and a sgRNA recognizing the donor recognition sequence (DRS), enabling cleavage and release of the donor payload for insertion and endogenous protein tagging with a spaghetti monster V5 (smV5) tag. The construct was also adapted for insertion of GFP or TurboID. (B) Experimental paradigm: AAVs were systemically delivered via retro-orbital (R/O) injection into 1-month-old *NG2-Cre;SELECTIV* or 2-month-old *MOBP-iCre;SELECTIV* mice and allowed to incubate for 3 weeks before analysis. (C-F) Endogenous tagging in 1-month-old *NG2-Cre;SELECTIV* mice injected with AAV-CAP-B10-GOI-KI viruses. Examples include β-Actin tagged with GFP in two adjacent cells (AAV-CAP-B10-Actb-GFP-KI; C; scale bar, 50 μm), Myelin basic protein (MBP) tagged with smV5 in a single cell (AAV-CAP-B10-MBP-smV5-KI; D; scale bar, 20 μm), Neurofascin tagged with smV5 in a single cell (AAV-CAP-B10-Nfasc-smV5-KI; E; scale bar, 20 μm), and Ankyrin-G tagged with smV5 in three adjacent cells (AAV-CAP-B10-Ank3-smV5-KI; F; scale bar, 50 μm). (G-J) Endogenous tagging in 2-month-old *MOBP-iCre;SELECTIV* mice. Examples include MBP tagged with smV5 in a single cell (AAV-CAP-B10-MBP-smV5-KI; G; scale bar, 20 μm) and Claudin-11 tagged with smV5 in a single cell (AAV-CAP-B10-Cldn11-smV5-KI; H; scale bar, 10 μm), showing paranodal colocalization with Caspr (magenta; I-J; scale bar, 10 μm). (K) MBP tagged with TurboID in 2-month-old *MOBP-iCre;SELECTIV* mice, with biotinylated proteins detected by streptavidin-488 (green) co-labeled with Caspr (magenta; scale bar, 20 μm). (L-N) Neurofascin tagged with TurboID in 1-month-old *NG2-Cre;SELECTIV* mice, showing paranodal localization of biotinylated proteins labeled with streptavidin-488 (green) that colocalize with Caspr (magenta; scale bar, 10 μm).

As a proof-of-principle, we applied the same KI constructs to OLs and successfully tagged multiple classes of proteins with distinct subcellular localizations. In *NG2-Cre;SELECTIV* mice, we achieved robust KI of β-Actin (cytoskeleton, **Fig. 4C**), MBP (myelin, **Fig. 4D**), AnkyrinG (paranodal scaffold protein, **Fig. 4E**), and Neurofascin (paranodal adhesion molecule, **Fig. 4F**). Remarkably, a single OL formed up to 70 paranodes with distinct axons, underscoring the extensive capacity of individual OLs to interact with multiple axons. GFP KI signals were detectable without antibody amplification, while smV5 staining yielded strong, unambiguous signal with a high signal-to-noise ratio. Due to the stochastic nature of NHEJ-based repair, KI events were sparse, which facilitated clear single-cell visualization against unlabeled neighbors. In *MOBP-iCre;SELECTIV* mice, however, KI was observed only for Actin and MBP, but not for AnkyrinG or Neurofascin, suggesting stage-dependent differences in protein turnover (**Fig. 4G**). We also tagged Claudin-11, an OL-enriched tight junction protein, and visualized its spiral paranodal structure wrapping around the axons, indicated by co-labeling with Caspr (**Fig. 4H-J**).

To extend and further illustrate the flexibility of the OASIS method, we replaced smV5 with TurboID, enabling both localization and enzymatic labeling by the promiscuous biotin-ligase. Consistent with the smV5 KI results, in *NG2-Cre;SELECTIV* mice, TurboID was successfully fused to endogenous MBP, AnkyrinG, and Neurofascin (**Fig. 4L-N**), while in *MOBP-iCre;SELECTIV* mice only MBP-TurboID was validated (**Fig. 4K**). Importantly, the TurboID fusion proteins were correctly localized and remained enzymatically active in OLs, as indicated by fluorophore-conjugated streptavidin labeling and immunofluorescence staining of Caspr (**Fig. 4K, L**). These results established proof-of-principle for future endogenous proteinbased proximity proteomics to bring novel mechanistic insights into axon-myelin interactions. To our knowledge, these results are the first successful AAV-mediated genome editing and endogenous protein tagging in adult OLs in vivo, highlighting both the versatility and the technical boundaries of the OASIS approach.

## Discussion

Here, we show that overexpression of the AAV receptor in oligodendrocyte lineage cells is sufficient to drive efficient and selective transduction of oligodendrocytes while markedly reducing neuronal infection. Using retro-orbital delivery, we achieved broad and uniform targeting across the brain with both AAV-PHP.eB and AAV-CAP-B10 capsids. This strategy proved effective throughout the lineage, from OPCs to mature myelinating oligodendrocytes, when combined with appropriate Cre drivers. We further demonstrated the utility of the CRISPR/Cas9 knock-in system by tagging diverse types of endogenous proteins, including cytoskeleton, myelin, cell adhesion molecules, scaffolding proteins, and tight junction proteins. Importantly, we established the feasibility of fusing exogenous enzymes such as TurboID to endogenous proteins, enabling enzymatic labeling across OL-lineage cells and at defined subcellular domains such as the paranode. Detection of paranode-specific biotinylation provides strong proof-of-principle for unbiased discovery of axon-myelin interactions through proximity proteomics. Collectively, our system represents a rapid and versatile alternative to conventional transgenic mouse models, allowing protein visualization, modification, and mechanistic investigation at single-cell resolution without reliance on commercial antibodies.

OASIS offers several advantages over conventional methods. First, it dramatically shortens the timeline and reduces the resources needed compared to generating and breeding multiple transgenic mouse lines, which typically takes months to years. In contrast, by designing gRNAs using existing databases and tools like CRISPOR^12^, using a streamlined universal cloning pipeline, in-house AAV packaging and quick systemic delivery, our approach only requires a few weeks from start to finish. This enables higher throughput testing of multiple candidate genes in parallel. Second, by introducing a universal smV5 epitope tag, we amplify signal intensity while avoiding nonspecific binding of poorly validated antibodies. Third, by targeting endogenous proteins in their native environment, we avoid potential artifacts introduced by overexpressing foreign cDNAs. Lastly, the inherently low efficiency of NHEJ repair enables sparse labeling, which allows clear subcellular visualization without dissociating individual cells in an in vitro culture system, preserving extracellular matrix and intercellular interactions critical for myelin biology.

Despite its utility, OASIS has several limitations. First, the choice of Cre drivers is critical. Interestingly, we achieved robust knock-in with *NG2-Cre; SELECTIV* mice but not *MOBP-iCre;SELECTIV* mice for certain paranodal proteins, despite both lines clearly showing efficient transduction by AAVs encoding CAG-flex-GFP. This may reflect the extremely long half-lives of some proteins that are rarely turned over during normal physiological conditions. For example, both AnkyrinG and Neurofascin are extremely stable ^13^. Therefore, successful DNA-level manipulations may not be able to translate into protein products in these contexts. Second, as with all CRISPR/Cas9based methods, off-target effects remain a risk. Therefore, multiple gRNAs should be used, and conclusions should be drawn only when identical localization patterns are observed across different targeting constructs. Third, knock-ins at extreme Nor C-terminal domains may disrupt localization or interactions of the innate protein. In the cases where either terminus of the target protein carries important localization motifs, intronic targeting should be selected instead. Finally, we caution that the absence of knock-in signal should not be interpreted as absence of the protein, since false negatives are more likely than false positives based on the design of the OASIS system, although a positive signal strongly confirms the presence of the target.

We anticipate that OASIS can be extended beyond knock-in of endogenous proteins. As a proof of principle, we demonstrated expression of GFP and efficient endogenous tagging through NHEJ-mediated knock-in, a relatively low-efficiency process compared to knock-out. Thus, the same framework may be adapted for CRISPR-based loss- or gain-of-function studies, although over-expression may be limited by AAV packaging capacity. To circumvent this, minimal functional motifs fused to reporters or enzymes may be used to enable precise domain targeting. For instance, the FIGQY motif of Neurofascin^14^ could be introduced together with exogenous TurboID to target the biotin-ligase to paranodes. OASIS could also be combined with complementary CRISPR technologies such as Hide-and-Seek^15^, which simultaneously deletes a gene while visualizing structural consequences. We anticipate the OASIS system could be extended to the study of not only OL-lineage cells, but other glial cells or specific subtypes of neurons. The stochastic, sparse labeling inherent to our system could also provide a natural platform for visualization of different sub-populations of cells. By “mixing-and-matching” AAVs with different fluorophores, one could also create an OL-specific “brainbow” system for lineage and connectivity studies.

Overall, OASIS offers a rapid, versatile, and generalizable platform for endogenous protein tagging in oligodendrocytes. By overcoming key limitations of antibody-based and transgenic approaches, OASIS provides a scalable platform for dissecting the molecular composition and dynamics of subcellular domains, such as the axon-glia interactions at paranodes. Beyond the oligodendrocyte lineage, this strategy can be readily adapted to other glial or neuronal populations, accelerating both mechanistic discovery and translational advances across the nervous system.

## Acknowledgements

This work was supported by a National Multiple Sclerosis Society Career Transition Fellowship to X.D. (NMSS-TA-2403-42997), NIH grants to M.N.R. (R35NS122073 and R01MH121544), the Adelson Medical Research Foundation (to M.N.R. and E.P.), and the US-Israel Binational Science Foundation (to M.N.R. and E.P.). We thank Dr. Yuki Ogawa and Dr. Yudong Gao for technical assistance and valuable input, and all members of the Rasband laboratory for helpful discussions.

## Author contributions

X.D.: Conceptualization, Methodology, Investigation, Formal analysis, Funding acquisition, Writing - original draft.

J.R.C.: Investigation, Formal analysis. Y.X.: Resources.

Y.W.: Formal analysis.

E.P.: Conceptualization, Funding acquisition.

M.N.R.: Conceptualization, Methodology, Supervision, Funding acquisition, Writing - review & editing.

## Materials and Methods

All experiments were conducted in accordance with protocols approved by the Institutional Biosafety Committees (IBC: D-303) and the Institutional Animal Care and Use Committee (IACUC: AN-4634) at Baylor College of Medicine, following the National Institutes of Health Guide for the Care and Use of Laboratory Animals.

### Animals

Mice were housed in groups under a 12 h light/12 h dark cycle at 22 ^°^C and 40-60% humidity, with ad libitum access to food and water. Health was monitored daily by veterinary staff at the Baylor College of Medicine Center for Comparative Medicine.

The “SELECTIV” mice (B6(129)-Igs2^tm1(CAG-AU040320/mCherry,-cas9*)Janc^/J; RRID:IMSR_JAX:037553) and NG2-Cre mice (B6;FVB-Ifi208^Tg(Cspg4-cre)1Akik^/J; RRID:IMSR_JAX:008533) were obtained from The Jackson Laboratory. Only female NG2-Cre mice were used for breeding to prevent germline transmission. Both sexes were used for experiments. Mobp-iCre and MOBP-iCreER^T2^ mice (RRID:MGI:8242146) were generously provided by Dr. Dwight Bergles. NG2-Cre SELECTIV, MOBP-iCre SELECTV, and MOBP-iCreER^T2^ SELECTIV mice were generated by breeding the SELECTIV mice with corresponding Cre lines. For MOBP-iCreER^T2^ SELECTIV mice, tamoxifen (20mg/ml in 95% corn oil and 5% ethanol) was administered intraperitoneally for five consecutive days (100mg/kg of body weight) to 8-week-old mice. One week later, titrated AAV was delivered systemically via retro-orbital (R/O) injection (100 µl per mouse).

Genotyping for all mice was carried out by standard PCR. The following primers were used for genotyping:

Cre Forward: 5’ - TGC TGT TTC ACT GGT TAT GCG G - 3’

Cre Reverse: 5’ - TTG CCC CTG TTT CAC TAT CCA G - 3’

MOBP-iCre Forward: 5’ - CCA AGC TCT GCA CCT TTG TT- 3’

MOBP-iCre Reverse: 5’ - CAG GTT TTG GTG CAC ACA GTC A - 3’

SELECTIV WT Forward: 5’ - AGT GGG ACT GCT TTT TCC AG - 3’

SELECTIV WT Reverse: 5’ - GAT CTG GGG CCA TAA ATG C - 3’

SELECTIV Mutant Forward: 5’ - GGC AAA CAC CTT TGA AGT CC - 3’

SELECTIV Mutant Reverse: 5’ - ACC TTC AGC TTG GCG GTC T - 3’

### Antibodies

Primary antibodies used for immunofluorescence included mouse monoclonal antibodies against Caspr (UC Davis/NIH NeuroMab Facility, Cat# K65/35, RRID:AB_2877274, 1:1000), NeuN (Millipore, Cat# MAB377, RRID:AB_2298772, 1:1000), Sox10 (Santa Cruz Biotechnology, Cat# sc-365692, RRID:AB_10844002, 1:200), and V5 (Innovative Research, Cat# R960CUS, RRID:AB_159298, 1:1000); rat monoclonal MBP (Abcam, Cat# ab7349, RRID:AB_305869, 1:1000); rabbit monoclonal V5 (Cell Signaling Technology, Cat# 13202, RRID:AB_2687461, 1:1000); rabbit polyclonal GFP (Invitrogen, Cat# A11122, RRID:AB_2307355, 1:1000), Caspr (Abcam, Cat# ab34151, RRID:AB_869934, 1:500), and Olig2 (Millipore, Cat# AB9610, RRID:AB_570666, 1:500); and chicken polyclonal GFP (Aves Labs, Cat# GFP-1020, RRID:AB_10000240, 1:1000). Secondary antibodies were obtained from Thermo Fisher Scientific and Jackson ImmunoResearch and, whenever possible, were chosen to match the isotype of the primary antibody.

### Immunofluorescence labeling

Mice were anesthetized with isoflurane and transcardially perfused with ice-cold PBS followed by 4% paraformaldehyde (PFA, pH 7.4). Brains were carefully dissected and post-fixed in 4% PFA at 4 ^°^C overnight. Tissues were then cryoprotected by sequential incubation in 20% and 30% sucrose in 0.1 M phosphate buffer (PB) for 24 h each at 4 ^°^C. Brains were embedded in OCT compound and equilibrated at -20 ^°^C before sectioning at 10-20 µm thickness with a cryostat (Thermo Scientific: HM525NX). Sections were directly mounted onto Superfrost Plus slides (VWR: 48311-703).

Slides were washed three times in PBS (5 min each) at room temperature (RT), permeabilized with PBST (PBS + 0.3% Triton X-100) for 5 min at RT, and blocked in blocking solution (10% normal goat serum in PBST) for 1 h at RT. Hydrophobic barriers were drawn using an ImmEdge Pen (Vector Laboratories: H-4000), and primary antibodies diluted in blocking buffer were applied and incubated overnight at 4 ^°^C in a humidity-controlled chamber. The following day, slides were washed three times with PBS at RT, and incubated with secondary antibodies diluted in blocking solution for 1 h at RT. Slides were then washed three times, and mounted with VECTASHIELD HardSet Antifade Mounting Medium (Vector Laboratories: H-1400-10). Images were acquired using a Zeiss Apotome.2 microscope.

### AAV production and titration

15-cm cell culture dishes were coated with poly-L-lysine (PLL) for 10 min at room temperature and rinsed three times with dPBS. Approximately 2.2 × 10^6^ low-passage HEK293T cells were seeded per dish in complete medium (DMEM supplemented with 10% FBS and 1% penicillin/streptomycin) and cultured at 37 ^°^C with 5% CO_2_. After 24 h, cells were transfected with three plasmids: a serotype-specific capsid (AAV-PHP.eB or AAV-CAP-B10), a pHelper, and a gRNA-containing targeting vector, using PEI in OptiMEM. 24 h later, media was replaced with fresh complete medium, and cells were maintained for another 48∼72 h to allow AAV production. Both supernatant and cells were harvested as previously described (Yuki’s PNAS).

Briefly, supernatant was incubated with PEG-8000 for ≥3 h at 4 ^°^C and centrifuged at 10,000 × g for 15 min at 4 ^°^C, while AAV-producing cells were pelleted at 3,000 × g for 5 min at room temperature and lysed to extract viral particles. Viral fractions from both supernatant and cell pellets were combined, passed through a 0.45 µm PVDF syringe filter, concentrated using Amicon Ultra-15 centrifugal filters, and washed at least five times by centrifugation at 5,000 × g for ≥30 min each at 4 ^°^C. The final concentrated AAV preparation (∼100 µl per batch) was titrated by qPCR using the AAVPro Titration Kit (Takara).

### CRISPR/Cas9 genome-editing

Guide RNAs (gRNAs) targeting candidate genes were selected using CRISPOR (RRID:SCR_015935) or based on validated sequences from prior publications. Two gRNAs were designed per gene whenever possible. gRNA sequences used for endogenous tagging were:

mActb gRNA: 5’-CGCAGCGATATCGTCATCCA-3’

mAnk3 gRNA: 5’- GAA GAA GGA AAT CCG GAA CGT GG -3’ ‘

mCldn11 gRNA#1: 5’- GGA GCA GCC CTC TTA GAC ATG GG -3’

mCldn11 gRNA#2: 5’- CAA GAG TGC CCA TGT CTA AGA GG -3’

mMBP gRNA#1: 5’- GGG CTC TCA GCG TCT CGC CAT GG -3’

mMBP gRNA#2: 5’- CAG CCG CTC TGG ATC TCC CAT GG -3’

mNfasc gRNA#1: 5’- TGG GAT ACT CCG TCA GGC AAG GG -3’

mNfasc gRNA#2: 5’- CCA TCT ATT CCC TTG CCT GAC GG -3’

### Image processing

ZEISS ZEN Microscopy Software (RRID:SCR_013672) was used to process images in batch. ImageJ (RRID:SCR_003070) was used for quantification of colocalization. Adobe Photoshop (RRID:SCR_014199) and Adobe Illustrator (RRID:SCR_010279) were used for generating figures.

## References

1. Rhodes, K. J. & Trimmer, J. S. Antibodies as valuable neuroscience research tools versus reagents of mass distraction. J Neurosci 26, 8017–8020, doi:10.1523/JNEUROSCI.2728-06.2006 (2006).

2. Suzuki, K. & Izpisua Belmonte, J. C. In vivo genome editing via the HITI method as a tool for gene therapy. Journal of human genetics 63, 157–164, doi:10.1038/s10038-017-0352-4 (2018).

3. Willems, J. et al. ORANGE: A CRISPR/Cas9-based genome editing toolbox for epitope tagging of endogenous proteins in neurons. PLoS Biol 18, e3000665, doi:10.1371/journal.pbio.3000665 (2020).

4. Gao, Y. et al. Plug-and-Play Protein Modification Using Homology-Independent Universal Genome Engineering. Neuron 103, 583–597 e588, doi:10.1016/j.neuron.2019.05.047 (2019).

5. Zhang, W. et al. A hierarchy of PDZ domain scaffolding proteins clusters the Kv1 K(+) channel protein complex at the axon initial segment. Sci Adv 11, eadv1281, doi:10.1126/sciadv.adv1281 (2025).

6. Zhang, W. et al. Immunoproximity biotinylation reveals the axon initial segment proteome. Nature communications 14, 8201, doi:10.1038/s41467-023-44015-2 (2023).

7. Ogawa, Y. et al. Antibody-directed extracellular proximity biotinylation reveals that Contactin-1 regulates axo-axonic innervation of axon initial segments. Nature communications 14, 6797, doi:10.1038/s41467-023-42273-8 (2023).

8. Zengel, J. et al. Hardwiring tissue-specific AAV transduction in mice through engineered receptor expression. Nature methods 20, 1070–1081, doi:10.1038/s41592-023-01896-x (2023).

9. Chan, K. Y. et al. Engineered AAVs for efficient noninvasive gene delivery to the central and peripheral nervous systems. Nat Neurosci 20, 1172–1179, doi:10.1038/nn.4593 (2017).

10. Stanton, A. C. et al. Systemic administration of novel engineered AAV capsids facilitates enhanced transgene expression in the macaque CNS. Med 4, 31–50 e38, doi:10.1016/j.medj.2022.11.002 (2023).

11. Goertsen, D. et al. AAV capsid variants with brain-wide transgene expression and decreased liver targeting after intravenous delivery in mouse and marmoset. Nat Neurosci 25, 106–115, doi:10.1038/s41593-021-00969-4 (2022).

12. Concordet, J. P. & Haeussler, M. CRISPOR: intuitive guide selection for CRISPR/Cas9 genome editing experiments and screens. Nucleic Acids Res 46, W242–W245, doi:10.1093/nar/gky354 (2018).

13. Saifetiarova, J., Taylor, A. M. & Bhat, M. A. Early and Late Loss of the Cytoskeletal Scaffolding Protein, Ankyrin G Reveals Its Role in Maturation and Maintenance of Nodes of Ranvier in Myelinated Axons. J Neurosci 37, 2524–2538, doi:10.1523/JNEUROSCI.2661-16.2017 (2017).

14. Tuvia, S., Garver, T. D. & Bennett, V. The phosphorylation state of the FIGQY tyrosine of neurofascin determines ankyrin-binding activity and patterns of cell segregation. Proc Natl Acad Sci U S A 94, 12957–12962 (1997).

15. Ogawa, Y., Nguyen, D. V. M., Ogawa, A. & Rasband, M. N. Hide- and-Seek genome editing reveals that Gephyrin is required for axoaxonic synapse assembly. Proc Natl Acad Sci U S A 122, e2500726122, doi:10.1073/pnas.2500726122 (2025).

